# Microbial biodeterioration of eighteenth-century oil paintings in Orosi, Costa Rica, and *in vitro* evaluation of volatile essential oil components as antimicrobials

**DOI:** 10.64898/2026.05.07.723565

**Authors:** Fabiola María Madrigal-Rodríguez, Priscilla Castro-Vargas, Daniela Jaikel-Víquez, Maríamalia Cob-Delgado, Roberto Marín-Delgado, José Alejandro Álvarez-Quesada, Mario Cubero-Campos, Montserrat Jarquín-Cordero, Jesús A. Espinoza-Valverde, Óscar A. Herrera-Sancho, Mauricio Redondo-Solano

## Abstract

Microbial colonization is a major cause of deterioration in paintings, leading to discoloration, pigment degradation, and loss of structural integrity. While biodeterioration of artworks has been studied in temperate climates, tropical environments remain underexplored despite their high humidity and temperature, which promote microbial growth. This study assessed the microbiological deterioration of two eighteenth-century oil paintings, *La Muerte de San José* and *Virgen de Guadalupe*, located in Orosi’s Colonial Church and Religious Art Museum, Costa Rica. Microorganisms were isolated and identified using VITEK® 2, microscopy, and MALDI-ToF analysis, and their biofilm-forming capacity was evaluated. Additionally, the antimicrobial activity of six essential oil components was tested using direct and indirect contact assays. Twenty-three bacterial species and fifteen fungal genera were identified, with *Bacillus*, *Staphylococcus*, *Cladosporium*, and *Aspergillus* among the most common. Notably, *La Virgen de Guadalupe* displayed the highest microbial diversity, reflected in a high Shannon index, indicative of a more complex microbial community. Several isolates displayed strong biofilm formation, particularly *Bacillus subtilis/amyloliquefaciens/vallismortis* and *Staphylococcus saprophyticus*. Linalool exhibited the strongest inhibitory activity, achieving complete bacterial growth inhibition in non-contact assays. Environmental monitoring revealed persistently elevated relative humidity and CO₂ levels during the study period. Together, these results reveal the complex microbial ecology of tropical heritage paintings and demonstrate that volatile essential oil components can serve as candidates for low-impact antimicrobial strategies in preventive conservation.

**Importance:** Understanding the microbiological deterioration of cultural heritage in tropical environments is crucial for designing sustainable conservation strategies. While microbial colonization of artworks has been widely studied in temperate regions, data from tropical climates remain limited despite inherently favorable conditions for microbial proliferation. This study integrates microbiological, environmental, and physicochemical analyses to characterize microbial communities colonizing eighteenth-century oil paintings in Orosi, Costa Rica. By combining microbial identification, biofilm quantification, and essential oil biocide testing, it bridges applied microbiology and cultural heritage conservation. The finding that volatile components such as linalool inhibit biofilm-forming bacteria without direct contact highlights their potential as eco-friendly, noninvasive antimicrobial alternatives to conventional biocides. These results expand the understanding of biodeterioration dynamics under tropical conditions and offer a practical framework for developing sustainable, evidence-based conservation protocols that protect both heritage materials and the environment.

**Graphical abstract:** Figure 0.
Artistic visualization of the geographical context of the studied artworks and the multidisciplinary analytical approaches applied, highlighting the diversity of microorganisms identified (illustration by Keylin Ureña-Alvarado).

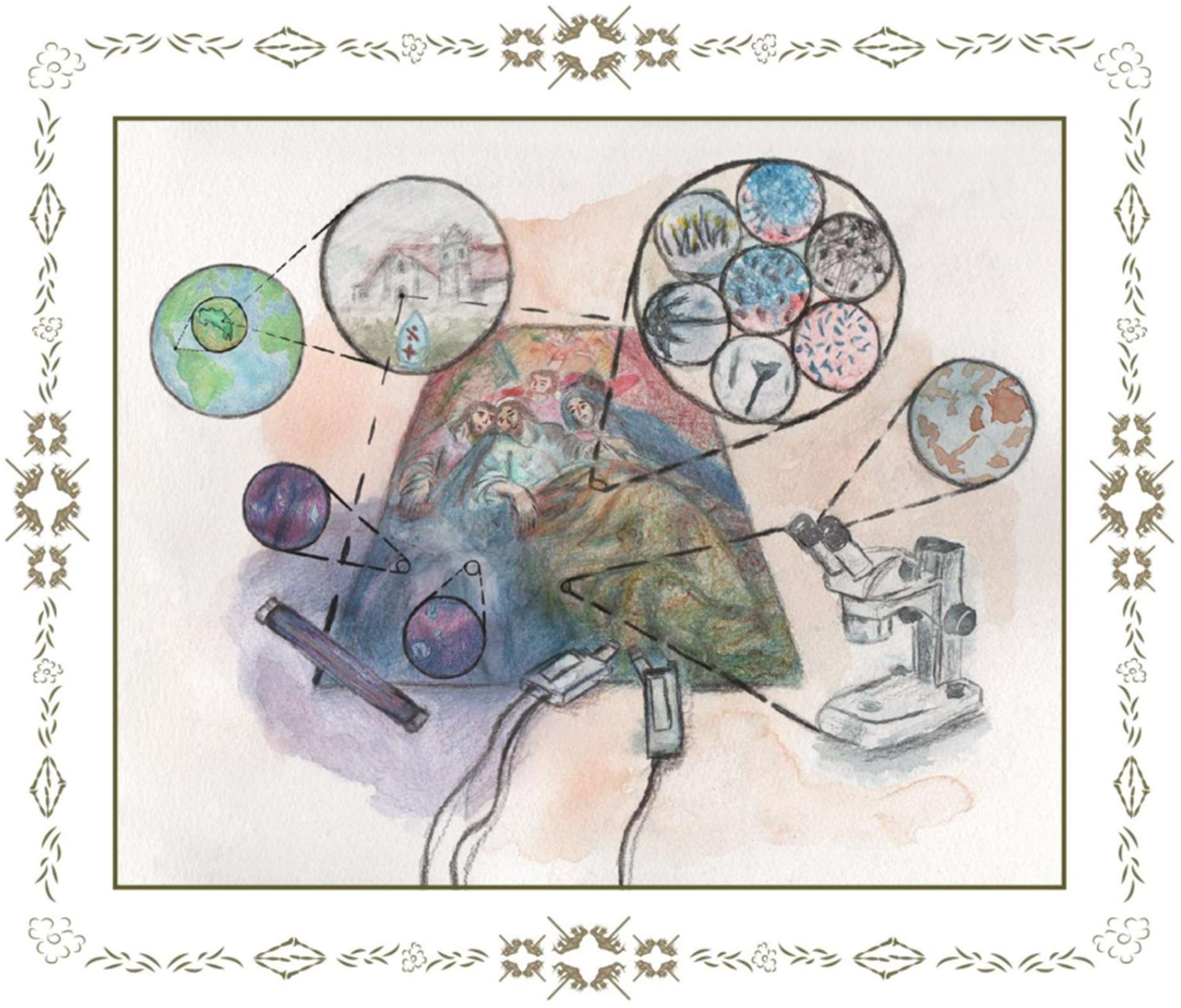

## Introduction

Cultural heritage represents a tangible record of human identity, artistic expression, and historical continuity (1, 2). However, the preservation of these materials is constantly challenged by physical, chemical, and biological deterioration processes. Among these, microbial biodeterioration has been recognized as a leading cause of irreversible damage to artworks, particularly in organic substrates such as canvas paintings, which provide nutrients and moisture favorable for microbial colonization (3, 4).

Microorganisms can colonize the surface and inner layers of artworks, producing pigments, organic acids, and extracellular enzymes that alter color, weaken fibers, and degrade binding media. The formation of biofilms further amplifies these effects by enhancing microbial adherence, nutrient retention, and resistance to environmental stress or cleaning agents (5, 6). Biofilm formation allows persistent deterioration over time, even under apparently controlled conditions, as metabolic activity continues within the polymeric matrix (6, 7).

While biodeterioration processes have been investigated in temperate and arid climates (8, 9), tropical environments remain understudied despite their constant high humidity and temperature, which favor microbial growth, sporulation, and biofilm development (10–12). The combination of high relative humidity and moderate temperatures typical of indoor heritage environments promotes microbial colonization and enzymatic activity on organic materials such as canvas and paint layers (1, 8, 13). Nevertheless, few studies have characterized the microbiological deterioration of cultural heritage in tropical Latin America, where historical artworks are often preserved in churches or museums lacking climate control.

Conventional conservation treatments rely on synthetic biocides such as quaternary ammonium compounds, alcohols, and chlorinated substances, which can effectively reduce microbial loads but also alter pigments, affect binder chemistry, or pose health and environmental risks (7, 14). Consequently, there is a growing movement toward sustainable, bio-based alternatives that combine antimicrobial efficacy with material compatibility (15).

Essential oils and their volatile components, such as linalool, α-terpineol, and eucalyptol, have demonstrated strong antifungal and antibacterial activities, making them promising candidates for cultural heritage applications (16–18). The antimicrobial activity of essential oils can occur through their volatile fractions, which diffuse in the vapor phase and inhibit microbial growth without requiring direct contact with the artwork surface; this has been demonstrated on stone, mural painting, and canvas substrates (14, 15).

In the present study, the microbiological deterioration of two eighteenth-century oil paintings, *La Muerte de San José* and *Virgen de Guadalupe*, from the Colonial Church and Religious Art Museum of Orosi, Costa Rica, was investigated. The culturable microbial communities were characterized, and their biofilm-forming capacity was evaluated. Environmental parameters, including temperature, humidity, and CO₂ concentration, were monitored as potential drivers of microbial colonization and community complexity, as reflected by high Shannon diversity index values. Furthermore, the antimicrobial activity of selected essential oil components was analyzed through both direct and indirect contact assays. By integrating microbiological, environmental, and chemical approaches, this research provides new insights into biodeterioration processes under tropical conditions and supports the development of eco-friendly, volatile-based strategies for the preventive conservation of cultural heritage materials.

## Materials and Methods

### Study site and sampling periods

Sampling and experimental procedures were performed between August and December 2022. Samples were obtained from the oil painting *La Muerte de San José* located in the Religious Art Museum in Orosi, Cartago, Costa Rica, and from the oil painting *Virgen de Guadalupe* located in the Colonial Church in Orosi, Cartago, Costa Rica (locations shown in Fig. S1). Samples were processed at the Laboratory for Research and Training in Food Microbiology (LIMA), the Mycology Laboratory, and the Research Center for Tropical Diseases (CIET), School of Microbiology, Universidad de Costa Rica.

### Sampling of paintings

The methodology described by Rivera-Romero *et al.* (19) was used with minor modifications. Ultraviolet and visible light photographs (Fig. 1) were used to select sampling sites. Samples were taken by swabbing the surface of selected areas of interest in different directions using a dry sterile cotton swab. A total of 24 sampling sites were selected on each painting, and each was sampled separately; a control zone with no visible damage according to photographic documentation was also selected for each painting. After sampling, the cotton swab was placed inside a tube with 5 mL of 0.1% sterile peptone water (PW). Sampling sites are shown in Fig. 1 and sampling codes in Table S1.

**Figure 1.**
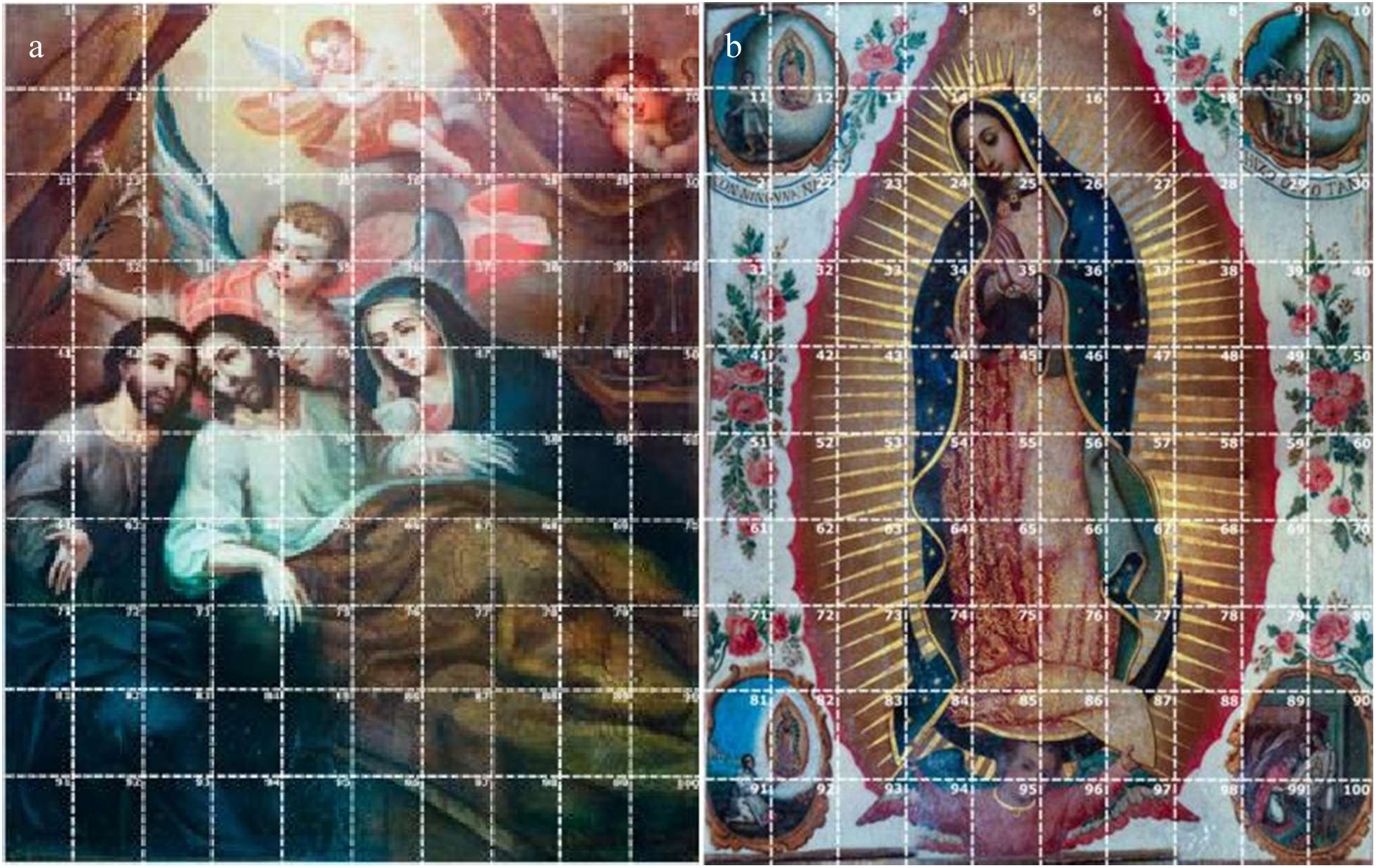
Sampling grid defined for the pictorial works under study. (a) *La Muerte de San José*, oil painting attributed to Cristóbal de Villalpando (18th century). (b) *Virgen de Guadalupe*, oil painting attributed to Miguel Cabrera (18th century). Photographs courtesy of Sandí-Delgado (2022).

### Isolation and purification of microbial colonies

For the isolation and purification of bacterial and fungal colonies, the methodology described by Rivera-Romero *et al.* (19) was used with some modifications. The suspensions were mixed and streaked onto blood agar (BA) and Sabouraud glucose agar (SGA) using the original sampling swab. BA plates were incubated at 35 °C for 48 h. Colonies of interest were subcultured on BA and incubated at 35 °C for 24 h. This procedure was repeated until purified bacterial strains were obtained, which were then streaked onto Trypticase soy agar (TSA) plates.

SGA plates were incubated at room temperature for three weeks and checked daily. Subcultures of distinct morphotypes were performed on SGA plates supplemented with antibiotics and on SGA or potato dextrose agar slants. This procedure was repeated when different morphotypes were observed on the same plate, until isolated strains were obtained. Once isolated, subcultures were performed to obtain two tubes for each strain.

### Microbial identification

Microbial identification followed Rivera-Romero *et al.* (19) with minor modifications. Bacterial colonies were initially characterized by Gram staining and then identified using the VITEK® 2 system. Bacterial isolates were stored at –80 °C in Brain Heart Infusion broth supplemented with 20% glycerol.

Initial identification of molds was performed through microscopic observations using lactophenol for dematiaceous (black) molds and lactophenol cotton blue for hyaline molds. Molds were classified using dichotomous keys based on morphology and spore type. To promote sporulation, non-sporulating molds were subcultured on “Lactrimiel” medium (honey, powdered milk and oats) and on oat agar. In some cases, VITEK® 2 was used for yeast identification, and final mold identification was performed by matrix-assisted laser desorption ionization–time of flight mass spectrometry (MALDI-ToF-MS), using the MBT Biotyper system (Bruker Daltonics). Spectra were analyzed using the Bruker Library and the Mass Spectra Identification platform (https://msi.happy-dev.fr).

### Characterization of bacterial biofilms

Biofilm formation was evaluated using the microtiter-plate method described by Carrillo-Zeledón *et al.* (20) and Stepanović *et al.* (21), with minor modifications. Twelve isolated and identified strains were selected (Table 1). Some strains were chosen because the same species were present on both paintings, others overlapped with species previously isolated from the paintings of the National Theatre of Costa Rica (NTCR) and preserved in the LIMA culture collection; or had been reported from cultural heritage elsewhere. Only strains with ≥85% confidence in VITEK® 2 identifications were included.

**Table 1.**
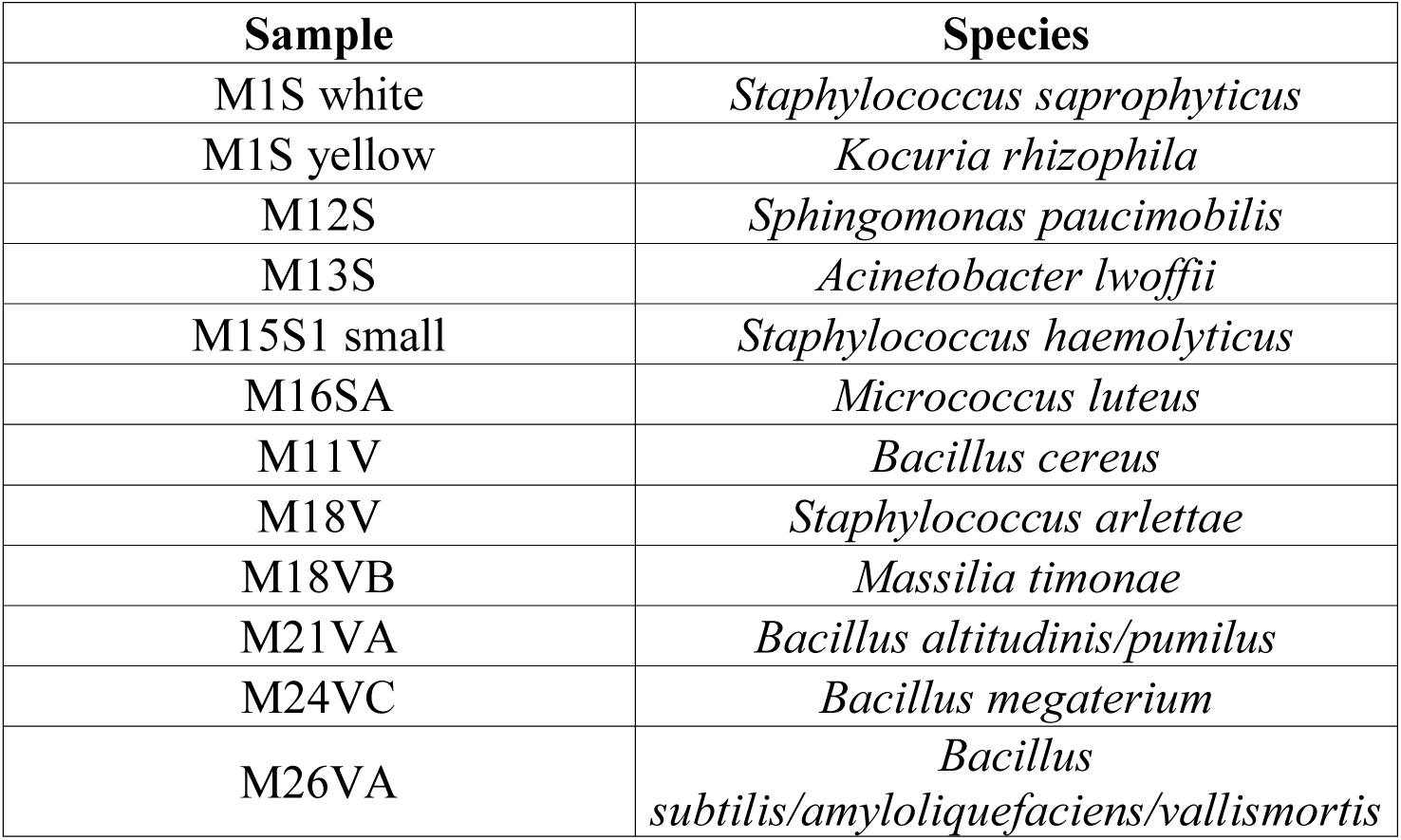
Selected strains for the characterization of bacterial biofilms.

| Sample | Species |
| --- | --- |
| M1S white | *Staphylococcus saprophyticus* |
| M1S yellow | *Kocuria rhizophila* |
| M12S | *Sphingomonas paucimobilis* |
| M13S | *Acinetobacter lwoffii* |
| M15S1 small | *Staphylococcus haemolyticus* |
| M16SA | *Micrococcus luteus* |
| M11V | *Bacillus cereus* |
| M18V | *Staphylococcus arlettae* |
| M18VB | *Massilia timonae* |
| M21VA | *Bacillus altitudinis/pumilus* |
| M24VC | *Bacillus megaterium* |
| M26VA | *Bacillus subtilis/amyloliquefaciens/vallismortis* |

For biofilm characterization, the selected strains were initially streaked on BA and incubated at 35°C for 24 h to verify purity and viability. Strains were then streaked on TSA and incubated at 35 °C for 24 h. Each bacterial suspension was adjusted to a McFarland 1 standard in 0.1% PW. Each well of a sterile 96-well microtiter plate was inoculated with 230 µL of Trypticase Soy Broth (TSB) and 20 µL of bacterial suspension. Negative controls contained only TSB. Each assay was performed in triplicate, and plates were incubated in a humid chamber at room temperature for 24 h.

After incubation, the contents of each well were discarded, and wells were washed three times with 300 µL of sterile water. Then, 250 µL of methanol was added to each well and left for 15 min to fix the cells. The remaining methanol was discarded, and the plates were air-dried. Wells were stained with 250 µL of crystal violet for 5 min, excess dye was discarded, and the wells were washed twice with 250 µL of sterile water. Finally, 250 µL of 33 % glacial acetic acid was added to solubilize the bound dye. Biofilm formation was quantified by measuring the optical density (OD) at 620 nm.

Classification of biofilm formation capacity followed the criteria defined by Stepanović *et al.* (21) and Stepanović *et al.* (22). A cutoff optical density (ODc) value was established as the mean OD of the negative controls plus three standard deviations. Isolates were classified as non-adherent, weak, moderate, or strong biofilm producers according to OD ranges (Table 2).

**Table 2.** Categories for the classification of biofilm formation capacity.

| Classification of biofilms forming capacity | Range of values |
| --- | --- |
| Non-adherent | $OD \leq OD_c$ |
| Weak biofilm producer | $OD_c < OD \leq (2 \times OD_c)$ |
| Moderate biofilm producer | $(2 \times OD_c) < OD \leq (4 \times OD_c)$ |
| Strong biofilm producer | $OD > (4 \times OD_c)$ |

### Biocide tests

The antimicrobial capacity of essential oils was evaluated using the methodology described by Gatti *et al.* (15) with some modifications. Three bacterial strains with strong or moderate biofilm-forming capacity and representing both paintings were selected (Table 3).

**Table 3.** Strains selected for the biocide tests.

| Sample | Species |
| --- | --- |
| M1S white | *Staphylococcus saprophyticus* |
| M1S yellow | *Kocuria rhizophila* |
| M26VA | *Bacillus subtilis/amyloliquefaciens/vallismortis* |

The selected essential oil components were eucalyptol, α-pinene, limonene, linalool, terpineol, and α-terpineol. Biocidal activity was evaluated in direct and indirect contact assays with the bacterial isolates. For direct contact, three BA plates per component were inoculated with 200 µL of a bacterial suspension (McFarland 1). Sterile filter paper disks impregnated with 50 µL of each component were placed on the inoculated plates and incubated at room temperature for 24 h. Negative and positive controls consisted of disks impregnated with sterile water and isopropanol, respectively. After incubation, inhibition zone diameters were measured.

For the non-contact test, BA plates were inoculated as described above, but 100 µL of each component was deposited on the inner surface of the plate lid. Plates were incubated inverted to prevent direct contact between the essential oil and the bacterial inoculum. After 24 h, bacterial growth inhibition was visually assessed, and inhibition diameters were measured when applicable.

### Environmental conditions

The meteorological station used in this study has been described previously (23). The station was placed in the same room as the artworks under study for approximately 8 days starting on March 21, 2021. The meteorological variables recorded during this period were temperature, relative humidity, and carbon dioxide (CO_2_) concentration. To ensure correct operation and avoid misreadings, the station was checked daily by one of the researchers.

### X-ray fluorescence spectroscopy (XRF)

XRF measurements were performed *in situ* using an Amptek Mini-X X-ray tube with a silver target and a 2-mm diameter collimator, providing a 5° X-Ray cone. To increase measurement reliability, the X-ray beam axis was set at an angle of 67.5° with respect to the plane of each painting. A silicon PIN X-ray detector (Amptek XR-100CR) was positioned at 45° to the primary beam and connected to an Amptek PX4 pulse processor. Spectra were acquired using Amptek acquisition software. The system was calibrated using titanium and molybdenum plates by adjusting peak positions to their true energies. Operating conditions were 34.9 kV and 15.4 μA, providing an effective energy range of 1.8-13.3 keV and a maximum energy resolution of 270 eV. The detector head was fixed on a support and positioned approximately 1.5 cm from the surface, producing an analyzed spot of about 0.4 mm in diameter. The methodological approach followed Rivera-Romero *et al.* (24). Spectral analysis was performed with PyMca, and peaks were manually compared with reported reference spectra (25). Twenty points per painting were selected according to: (i) coverage of as many apparent colors as possible, (ii) reproducibility within color classes, and (iii) coverage of overall surface variability (Fig. S2 and Fig. S3).

### Diversity and Statistical Analysis

The methodology began with segmentation of each artwork image into a 10 × 10 grid of uniformly sized zones. Each zone was assigned a unique identifier to facilitate subsequent reference. Caution must be applied since only zones associated with microbiological sampling were retained for subsequent diversity and chromatic analysis. Microbiological data associated with the selected zones were processed to quantify community structure. Four complementary alpha diversity indices were computed: the Shannon–Wiener index (26), Simpson index (27), Chao1 estimator (28), and Fisher’s α (29), capturing richness, evenness, and sampling-adjusted richness. The Shannon–Wiener index emphasizes communities with high richness and uneven distributions, whereas the Simpson index highlights dominance by a few species. The Chao1 estimator provides an estimate of the total potential richness by accounting for unobserved taxa inferred from the number of rare species, and Fisher’s α is particularly useful for comparing samples with differing total abundances due to its low sensitivity to sampling effort. Beta diversity was quantified using the Bray–Curtis dissimilarity index (30), to assess compositional differences between communities and their relationship with the visual and conservation state of each zone.

These metrics were computed following standard ecological formulations. The Shannon index is defined as:

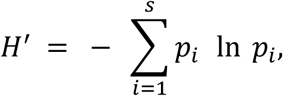

where *p*_i_ is the relative abundance of species *i* and *S* the observed richness. The Simpson index, expressed as 1 − *D* to represent diversity, is:

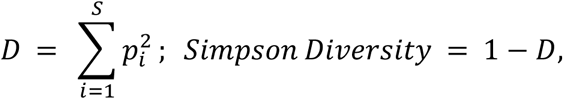

The Chao1 richness estimator incorporates information about rare taxa:

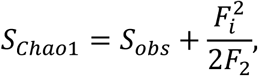

where *S*_obs_ is the observed richness, and *F₁* and *F₂* represent singleton and doubleton taxa, respectively. Fisher’s α was obtained from:

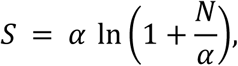

solved numerically for *α*, where *S* is species richness and *N* the total number of individuals.

The Bray–Curtis dissimilarity between zones *i* and *j* is computed as:

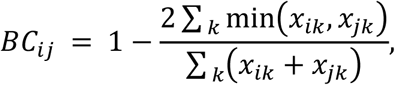

with *x*_ik_ and *x*_jk_ representing the abundance of species *k* in zones *i* and *j*.

After calculating diversity metrics, the chromatic characteristics of each microbiologically relevant zone are then extracted. Color features were quantified in the perceptually uniform CIELAB (LAB) color space (31). Each zone was summarized using normalized pixel-color histograms, capturing the frequency of hue-related and saturation-related components. Chromatic similarity between zones was assessed by pairwise comparison of these histograms using the Bhattacharyya distance (32):

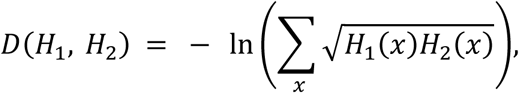

where *H*_1_ and *H*_2_ denote the normalized color histograms of two zones. The complete set of inter-zone chromatic distances was compiled into a distance matrix. Proximity profiles ordered by rank were generated for each zone, identifying its first and second-nearest chromatic neighbors. An annotated grid was generated, connecting each zone to its first and second nearest neighbors. To spatially display diversity information, a map of the Shannon contribution was overlaid. This map was constructed by superimposing two-dimensional Gaussian functions, where the amplitude of each Gaussian corresponds to the scalar contribution of the zone of interest to the overall Shannon index. This representation facilitates interpretation of potential spatial, ecological, and population-level relationships between microbial community structure and observed chromatic, and possibly pigment, patterns. All analyses were performed in Python 3.12 using the scikit-bio, scikit-learn, and pandas libraries.

## Results

### Microbial isolation and identification

#### Sampling overview

Sampling zones for each oil painting are presented in Fig. 2a and Fig. 3a. Sites used for microbial isolation are shown in pink, whereas control zones are shown in green (quadrants 26 and 15 for *La Muerte de San José* and *Virgen de Guadalupe*, respectively). Additional samples were taken from the back of the canvases and the wooden frames.

**Figure 2.**
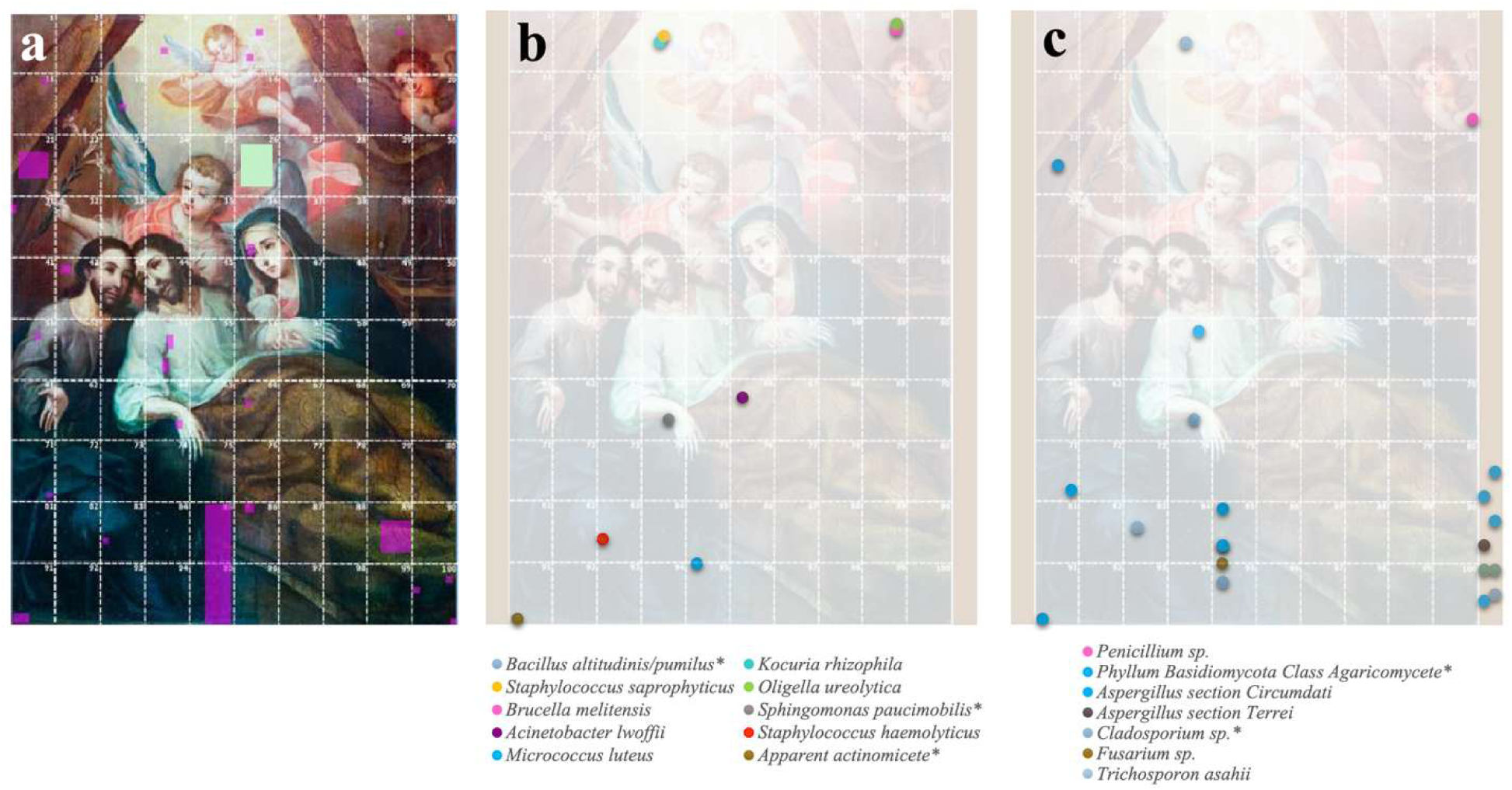
*La Muerte de San José*, oil painting attributed to Cristóbal de Villalpando (18th century), San José Museum of Religious Art, Orosi, Cartago, Costa Rica. (a) Visible-light image with the sampling grid; pink squares indicate sampled sites, and the green square indicates the control site. (b) Spatial distribution of isolated and identified bacterial species. (c) Spatial distribution of isolated and identified fungal taxa. Asterisks indicate taxa also detected on the back of the canvas.

**Figure 3.**
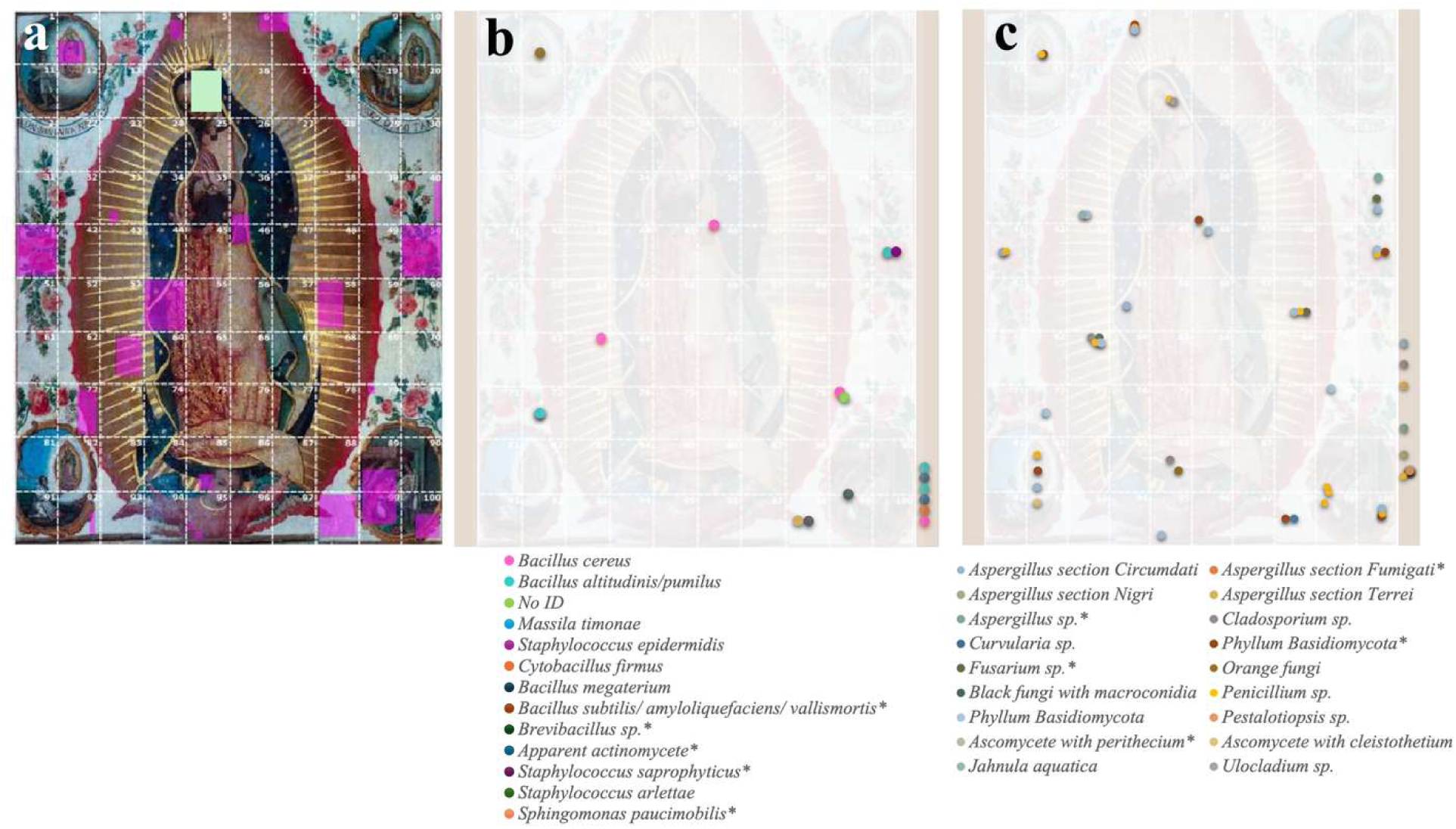
*Virgen de Guadalupe*, oil painting attributed to Miguel Cabrera (18th century), San José Colonial Church, Orosi, Cartago, Costa Rica. (a) Visible-light image with the sampling grid; pink squares indicate sampled sites, and the green square indicates the control site. (b) Spatial distribution of isolated and identified bacterial species. (c) Spatial distribution of isolated and identified fungal taxa. Asterisks indicate taxa also detected on the back of the canvas.

#### Bacterial isolation and identification

A total of 80 isolates and 23 bacterial species were obtained from both paintings. Nine species were identified from *La Muerte de San José*, including *Bacillus altitudinis/pumilus, Kocuria rhizophila, Staphylococcus saprophyticus, Oligella ureolytica*, *Brucella melitensis*, *Sphingomonas paucimobilis*, *Acinetobacter lwoffii*, *Micrococcus luteus* and *Staphylococcus haemolyticus.* Given the unexpected identification of *B. melitensis* by VITEK®, this result should be interpreted with caution, as environmental *Brucella*-like organisms may yield false-positive profiles in automated identification systems (33, 34). *B. altitudinis/pumilus* was isolated only from the back of the canvas (Fig. 2b). *S. paucimobilis* (n = 4) and *S. saprophyticus* (n = 2) were the most frequent isolates, while all other species were represented by a single isolate (n = 1).

In *Virgen de Guadalupe*, 11 species were identified, including *Bacillus cereus, B. altitudinis/pumilus, Massilia timonae, Staphylococcus epidermidis, Staphylococcus arlettae, Cytobacillus firmus, Bacillus megaterium*, *S. saprophyticus, Bacillus subtilis/amyloliquefaciens/vallismortis*, *Brevibacillus* sp., and *S. paucimobilis*. The last three were isolated only from the back of the canvas (Fig. 3b). The most frequent species were *B. altitudinis/pumilus* (n = 5), *B. cereus* (n = 4), *B. megaterium (*n = 2) and *S. saprophyticus* (n = 2); all other species were represented by a single isolate (n = 1). *B. altitudinis/pumilus*, *S. saprophyticus* and *S. paucimobilis* were recovered from both paintings.

### Fungal isolation and identification

A total of 155 fungal isolates were recovered from both paintings. Forty-three were retrieved from *La Muerte de San José*, among which eight genera were identified (Fig. 2c). In order of frequency: *Cylindrocarpon destructans* (n = 7), *Cladosporium* (n = 6; *C. cladosporioides* complex, *n* = 3, C. *tenuissimum*, *n* = 2, and *C. halotolerans, n* = 1; Fig. S4), *Aspergillus* (n = 5; *A. sclerotiorum n* = 4; *A.* section *Terrei, n* = 1), *Penicillium citrinum* (n = 3), *Fusarium* (n = 2; Fig. S4), *Coprinus* (n = 1), *Pestalotiopsis* (n = 1), and *Trichosporon asahii* (n = 1). Seventeen isolates could not be identified due to lack of sporulation and absence of matching reference material in the consulted databases, highlighting the limitations of morphology-based fungal identification.

The remaining 112 isolates were obtained from *Virgen de Guadalupe* (Fig. 3c), where 15 genera were identified. In order of frequency, *C. destructans* (n = 17), *Penicillium* (n = 13; Fig. S4), *Aspergillus* (n = 8; *A. scle*rotiorum, n = 2, *A.* section *Terrei*, n = 1; *A.* welwitschiae, n = 1; *A.* section *Fumigati*, n = 1; Fig. S4), *Cladosporium* (n = 5; *C. cladosporioides* complex, n = 4; *C. pseudocladosporioides*, n = 1), *Coprinus* (n = 7), *Fusarium* (n = 5), *Paecilomyces* (n = 3), *Geotrichum* (n = 2), *Hortaea* (n = 2), *Curvularia* (n = 1), *Jahnula aquatica* (n = 1), *Nakazawaea holstii* (n = 1), *Pestalotiopsis* (n = 1; Fig. S4), *T. asahii* (n = 1) and *Ulocladium* (n = 1).

### Bacterial biofilm formation

According to the classification of biofilm formation capacity, *S. saprophyticus* (OD_620nm_ = 0.298) and *B. subtilis/amyloliquefaciens/vallismortis* were classified as strong biofilm producers. In the case of *B. subtilis/amyloliquefaciens/vallismortis*, the high level of biofilm formation resulted in oversaturated OD readings; based on this qualitative observation, the strain was classified as a strong biofilm producer. *K. rhizophila* (OD_620nm_ = 0.149) was classified as a moderate biofilm producer. The remaining strains were classified as non-adherent, including *S. paucimobilis*, *A. lwoffii*, *S. haemolyticus*, *M. luteus*, *B. cereu*s, *S. arlettae*, *M. timonae*, *B. altitudinis/pumilus*, and *B. megaterium* (Table 4, Fig. 4).

**Figure 4.**
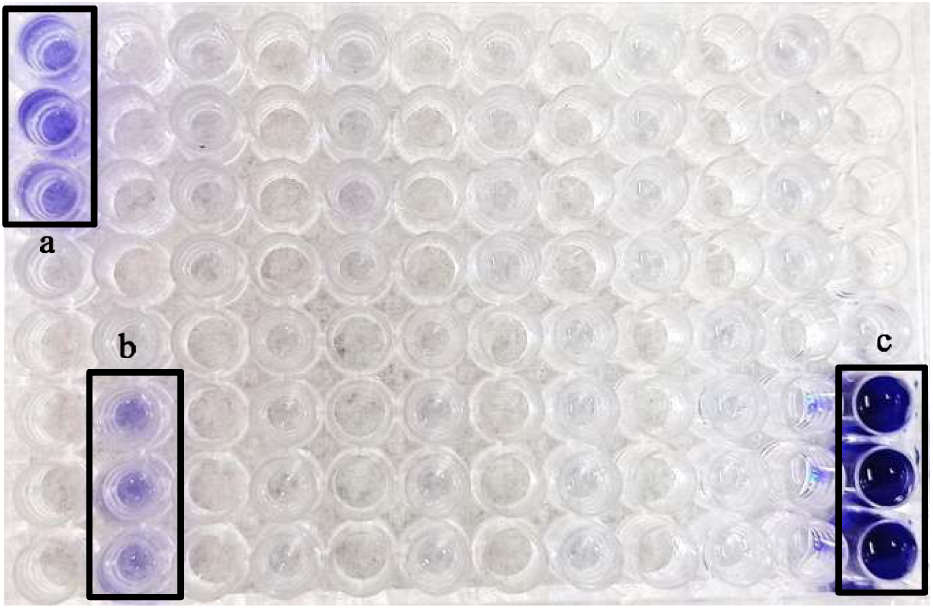
Microplate after crystal violet staining for biofilm detection and classification. (a) *S. saprophyticus*, strong biofilm producer; (b) *K. rhizophila*, moderate biofilm producer; (c) *B. subtilis/amyloquefaciens/vallismortis*, strong biofilm producer. Dark purple staining indicates higher biofilm biomass.

**Table 4.** Classification of strains by biofilm formation capacity.

| Strain | Average OD (620 nm) | Result |
| --- | --- | --- |
| *S. saprophyticus* (M1S white) | $0.298 \pm 0.036$ | Strong biofilm producer |
| *K. rhizophila* (M1S yellow) | $0.149 \pm 0.006$ | Moderate biofilm producer |
| *S. paucimobilis* (M12S) | $0.059 \pm 0.002$ | Non-adherent |
| *A. lwoffii* (M13S) | $0.058 \pm 0.002$ | Non-adherent |
| *S. haemolyticus* (M15S1 small) | $0.064 \pm 0.004$ | Non-adherent |
| *M. luteus* (M16SA) | $0.064 \pm 0.003$ | Non-adherent |
| *B. cereus* (M11V) | $0.059 \pm 0.002$ | Non-adherent |
| *S. arlettae* (M18V) | $0.062 \pm 0.003$ | Non-adherent |
| *B. altitudinis/pumilus* (M21VA) | $0.061 \pm 0.004$ | Non-adherent |
| *B. megaterium* (M24VC) | $0.062 \pm 0.002$ | Non-adherent |
| *B. subtilis/amyloliquefaciens/vallismortis* (M26VA) | Oversaturated | Strong biofilm producer |

### Antibacterial activity of essential oil components

In all cases, except for α-pinene against *B. subtilis//amyloliquefaciens/vallismortis*, inhibition zones were larger in non-contact assays than in direct-contact assays. Linalool produced the largest inhibition zones in non-contact assays for all three bacterial strains, achieving complete inhibition across the Petri dish (diameter 8.7 cm; Fig. 5). Accordingly, linalool exhibited the greatest antibacterial activity in this study (Table 5).

**Figure 5.**
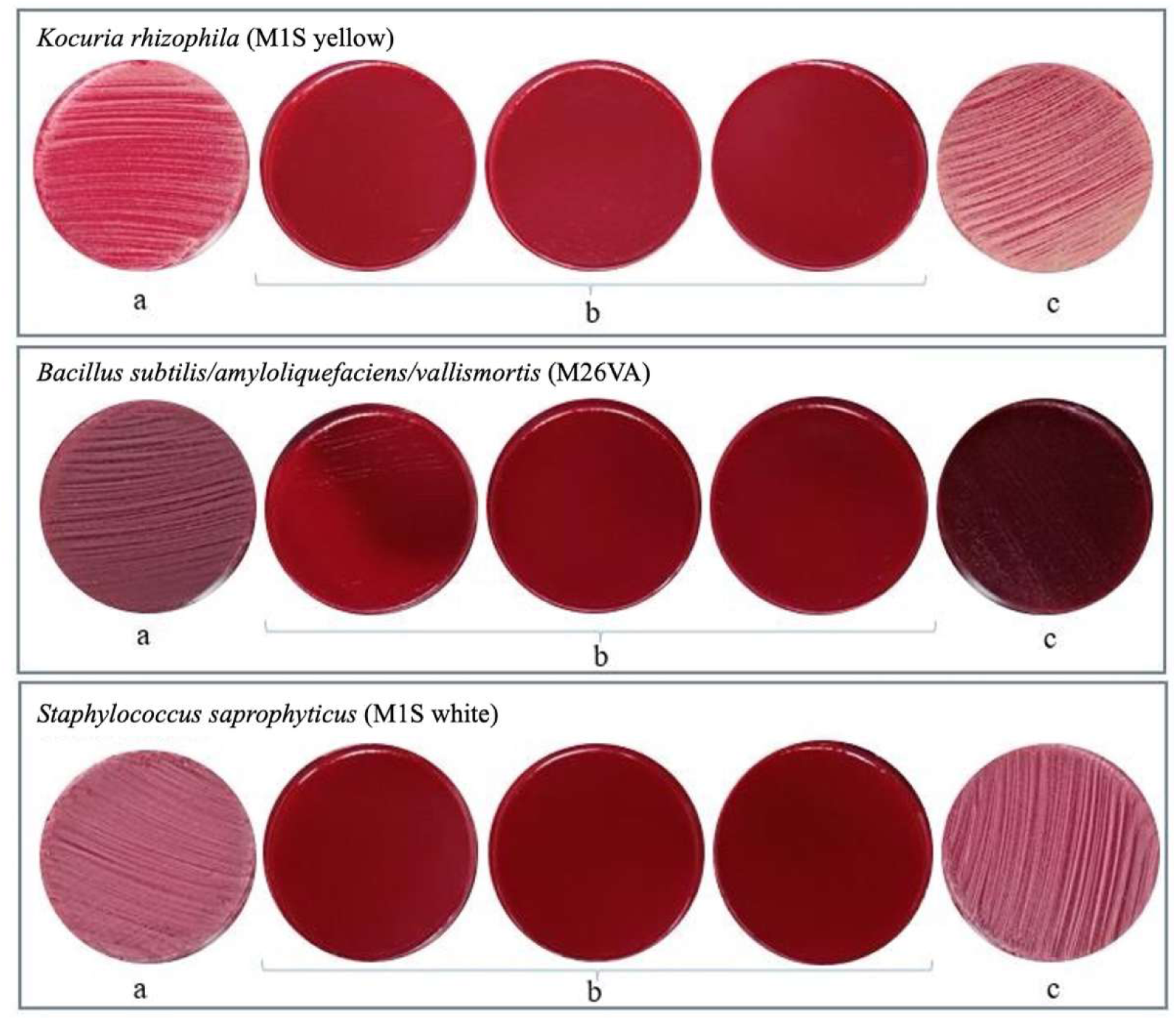
Representative results of indirect biocide assays for three bacterial isolates exposed to linalool. For each strain, panels show: (a) positive control (isopropanol), (b) plates treated with linalool (three replicates), and (c) negative control (no essential oil). Strains shown: *K. rhizophila* (M1S yellow), *B. subtilis/amyloliquefaciens/vallismortis* (M26VA), and *S. saprophyticus* (M1S white).

**Table 5.**
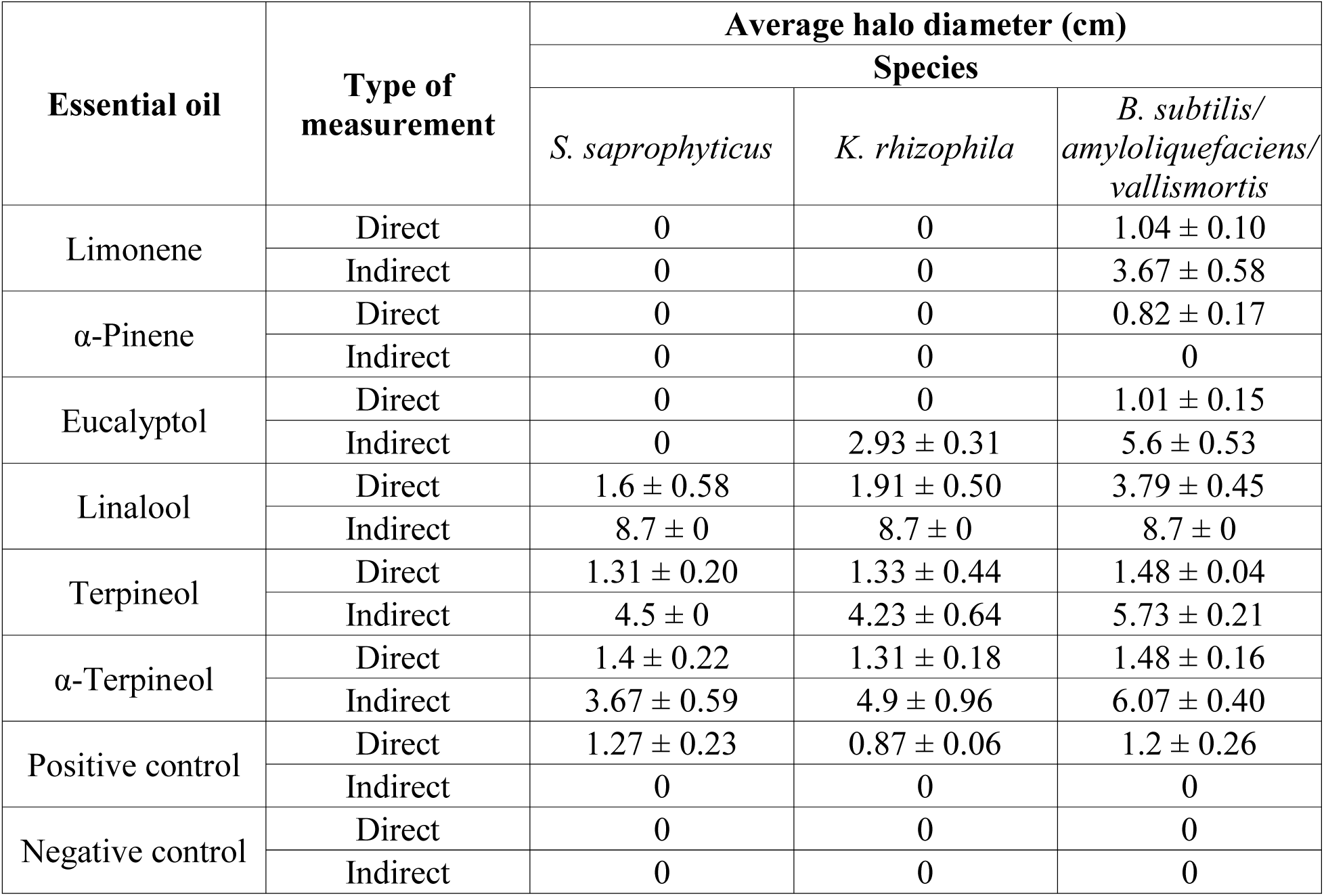
Inhibition zones (cm) produced by essential oil components in direct and indirect contact tests.

| Essential oil | Type of measurement | Average halo diameter (cm) |  |  |
| --- | --- | --- | --- | --- |
|  |  | Species |  |  |
|  |  | *S. saprophyticus* | *K. rhizophila* | *B. subtilis*/<br>*amyloliquefaciens*/<br>*vallismortis* |
| Limonene | Direct | 0 | 0 | 1.04 ± 0.10 |
| | Indirect | 0 | 0 | 3.67 ± 0.58 |
| α-Pinene | Direct | 0 | 0 | 0.82 ± 0.17 |
|  | Indirect | 0 | 0 | 0 |
| Eucalyptol | Direct | 0 | 0 | 1.01 ± 0.15 |
| | Indirect | 0 | 2.93 ± 0.31 | 5.6 ± 0.53 |
| Linalool | Direct | 1.6 ± 0.58 | 1.91 ± 0.50 | 3.79 ± 0.45 |
| | Indirect | 8.7 ± 0 | 8.7 ± 0 | 8.7 ± 0 |
| Terpineol | Direct | 1.31 ± 0.20 | 1.33 ± 0.44 | 1.48 ± 0.04 |
| | Indirect | 4.5 ± 0 | 4.23 ± 0.64 | 5.73 ± 0.21 |
| α-Terpineol | Direct | 1.4 ± 0.22 | 1.31 ± 0.18 | 1.48 ± 0.16 |
| | Indirect | 3.67 ± 0.59 | 4.9 ± 0.96 | 6.07 ± 0.40 |
| Positive control | Direct | 1.27 ± 0.23 | 0.87 ± 0.06 | 1.2 ± 0.26 |
|  | Indirect | 0 | 0 | 0 |
| Negative control | Direct | 0 | 0 | 0 |
|  | Indirect | 0 | 0 | 0 |

The second largest inhibition zone was observed for α-terpineol against *B. subtilis/amyloliquefaciens/vallismortis*, in the non-contact assay (6.07 ± 0.40 cm), followed by terpineol against *B. subtilis/amyloliquefaciens/vallismortis* (5.73 ± 0.21 cm) (Table 5). In direct-contact assays, the largest inhibition halo was observed for *B. subtilis/amyloliquefaciens/vallismortis* against linalool (3.79 ± 0.45 cm). However, these values were consistently smaller than those obtained in non-contact assays, where vapor phase exposure produced broader inhibition (Fig. 6).

**Figure 6.**
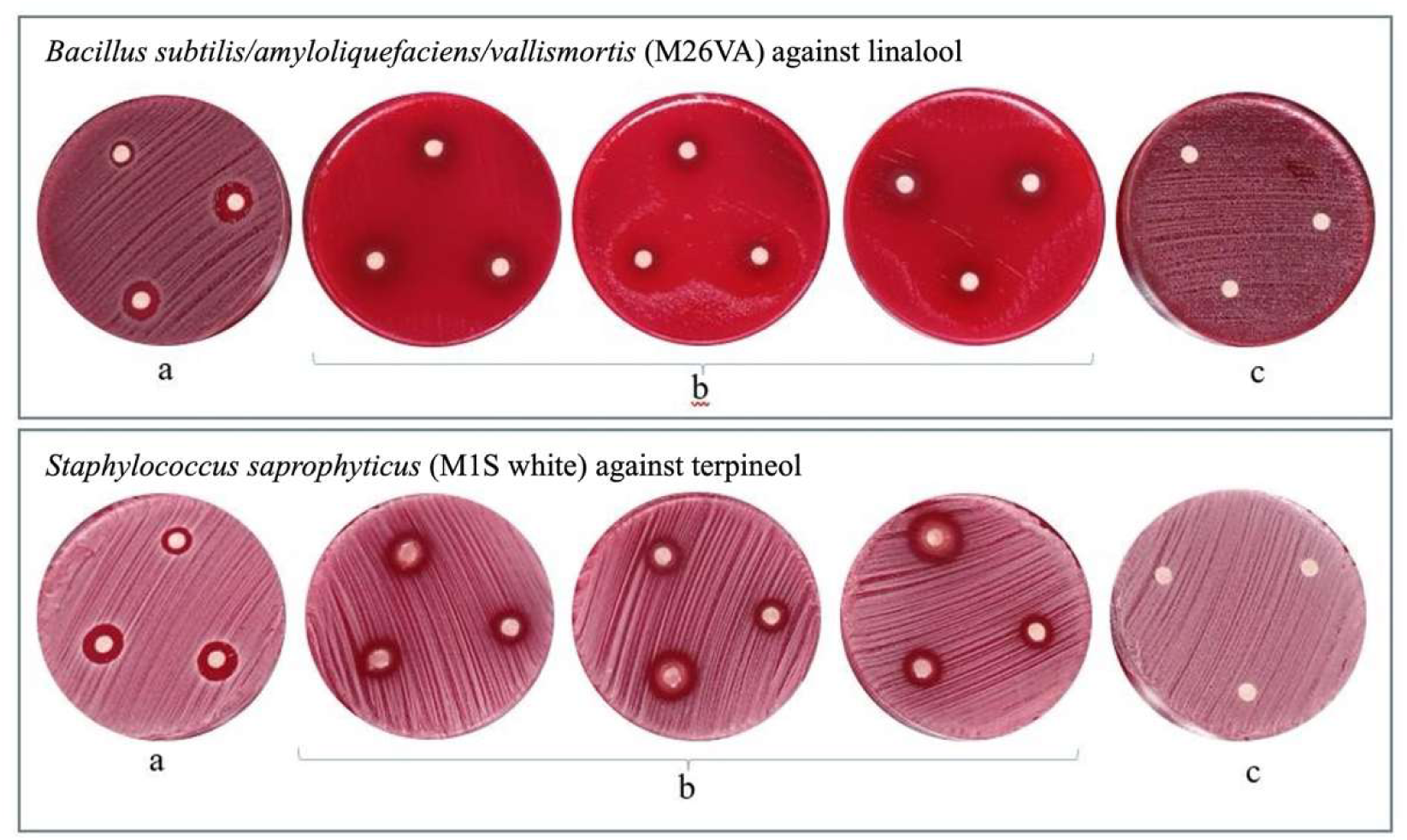
Direct-contact antibacterial assays with selected essential oil components. Top row: *Bacillus subtilis/amyloliquefaciens/vallismortis* (M26VA) exposed to linalool. Bottom row: *Staphylococcus saprophyticus* (M1S white) exposed to terpineol. (a) Positive controls (isopropanol); (b) plates with essential oil–impregnated disks and replicates; (c) negative controls (sterile water). Plates were incubated for 24 h at room temperature.

α-Pinene showed the lowest antibacterial activity, producing only a small inhibition zone (0.82 ± 0.17 cm) against *B. subtilis/amyloliquefaciens/vallismortis* in direct contact and no inhibition in other combinations (Table 5). Additionally, *S. saprophyticus* was the most resistant strain, not inhibited by limonene, α-pinene, or eucalyptol. *K. rhizophila* was also not inhibited by limonene or α-pinene. In contrast, *B. subtilis//amyloliquefaciens/vallismortis* was the most susceptible, being partially or completely inhibited in 91.67% of the tests and by all six components evaluated.

No negative control showed inhibition of bacterial growth or contamination of cultures or filter papers. Isopropanol (positive control) generated inhibition halos in direct-contact assays, but these were generally smaller than those produced by most essential oil components. Isopropanol had no detectable effect in non-contact assays, in contrast to several essential oil components that were active in the vapor phase.

### Environmental conditions

During the 8-day monitoring period, temperature ranged approximately from 23.5 to 26.0 °C, relative humidity from approximately 70 to 82%, and CO₂ concentration from approximately 300 to 410 ppm (Fig. 7). All three variables displayed marked daily oscillations, with temperature and CO₂ showing synchronous peaks during daytime hours, while relative humidity fluctuated inversely with temperature. These conditions indicate a persistently warm and highly humid microclimate favorable for microbial growth and biodeterioration processes.

**Figure 7.**
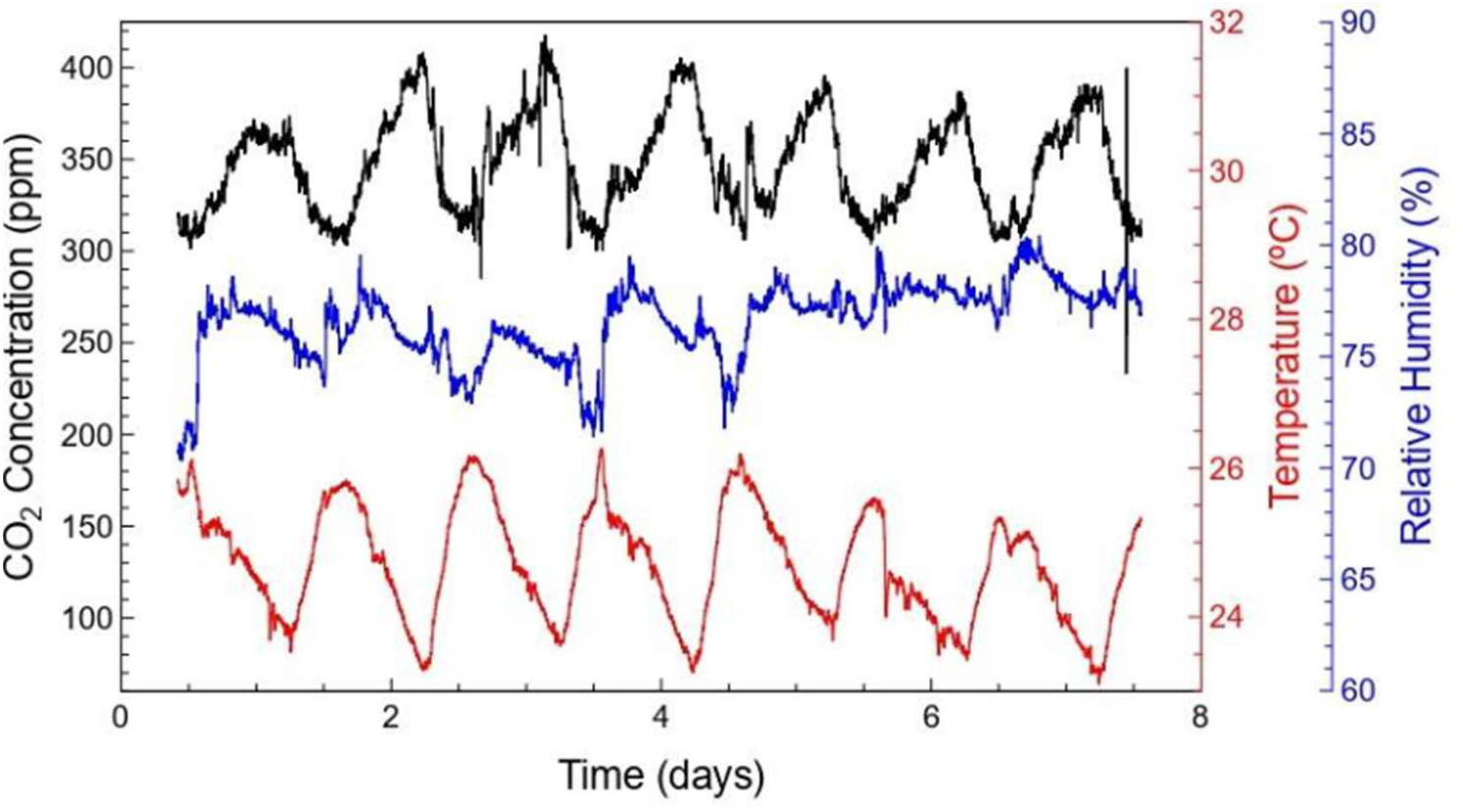
Time series of environmental parameters recorded every 2 min over 8 days in the exhibition space in Orosi, Cartago, Costa Rica. Temperature (red) ranged from approximately 23.5 to 26.0 °C, relative humidity (blue) from 70 to 82%, and CO₂ concentration (black) from 300 to 410 ppm.

### X-Ray fluorescence spectroscopy (XRF)

XRF measurements in *La Muerte de San José* suggested the presence of lead white, lead yellow, iron oxides, Prussian blue, and Scheele’s green. In *La Virgen de Guadalupe*, lead white, vermilion, iron oxides, Prussian blue, and Scheele’s green were detected. Specifically, the individual XRF spectra of white areas in backgrounds and clothing (VG11, VG29, VG91, MS46, MS52, MS54; see Fig. S2 and Fig. S3) displayed Pb lines, suggesting the presence of lead white (Pb₃(CO₃)₂(OH)₂). On the other hand, the background (VG29; see Fig. S3) showed Ti lines (TiO₂), which were inconsistent with the historical period (35), while for *La Muerte de San José* Fe lines were detected.

Fe lines in blue clothing and wings (VG35, VG44, VG57, VG84, MS13, MS72, MS82; see Fig. S2 and Fig. S3) may have indicated Prussian blue (Fe₄[Fe(CN)₆]₃). Red colors in the background, clothing, skin, wings, and fire (VG7, VG15, VG46, VG64, VG80, VG97, VG99, MS28, MS30; Fig. S2 and Fig. S3) showed Hg lines indicating vermilion (HgS). Yellow and brown areas in the background, clothing, hair, and a branch (VG5, VG89, MS1, MS4, MS5, MS9, MS25, MS31, MS44, MS68, MS86, MS90; Fig. S2 and Fig. S3) showed Fe lines consistent with ochre and sienna pigments (FeO(OH), FeO).

Background and leaf green areas (VG9, VG90, MS22; Fig. S2 and Fig. S3) displayed simultaneous Fe, Ca, and Cu lines; this suggested green earth or Prussian green with trace elements (Cu(OH)₂·((K(Mg,Fe²⁺)(Fe³⁺,Al)Si₄O₁₀)(OH)₂), ((K,Na)(Fe³⁺,Al,Mg)₂(Si,Al)₄O₁₀(OH)₂), Fe₄[Fe(CN)₆]₃). In addition, MS22 (Fig. S2) showed a peak corresponding to an As Kβ line; apart from the possible superposition of As Kα with Pb Lα lines (white background), this may have suggested Scheele’s green (Cu(AsO₂)₂). Gold lines were detected in VG77 (Fig. S3).

To determine the pigments present in the artworks, the measurements were correlated with historic data (36). Reference spectra were obtained from the Cultural Heritage Science Open Source (CHSOS) database.

### Diversity and Statistical Analysis

Across both artworks, the chromatic–microbiological overlays (Fig. 8) reveal subtle spatial associations between color similarity and microbial taxa distribution. In the figure, zones are connected according to chromatic proximity, defined by the closest (yellow) and second-closest (green) Bhattacharyya-distance relationships. Several localized patterns are observed in which chromatically similar zones tend to harbor overlapping or comparable microbial taxa. These associations suggest that surface properties indirectly reflected by color—such as pigment composition, surface texture, or micro-scale moisture retention—may contribute to structuring microbial colonization patterns.

**Figure 8.**
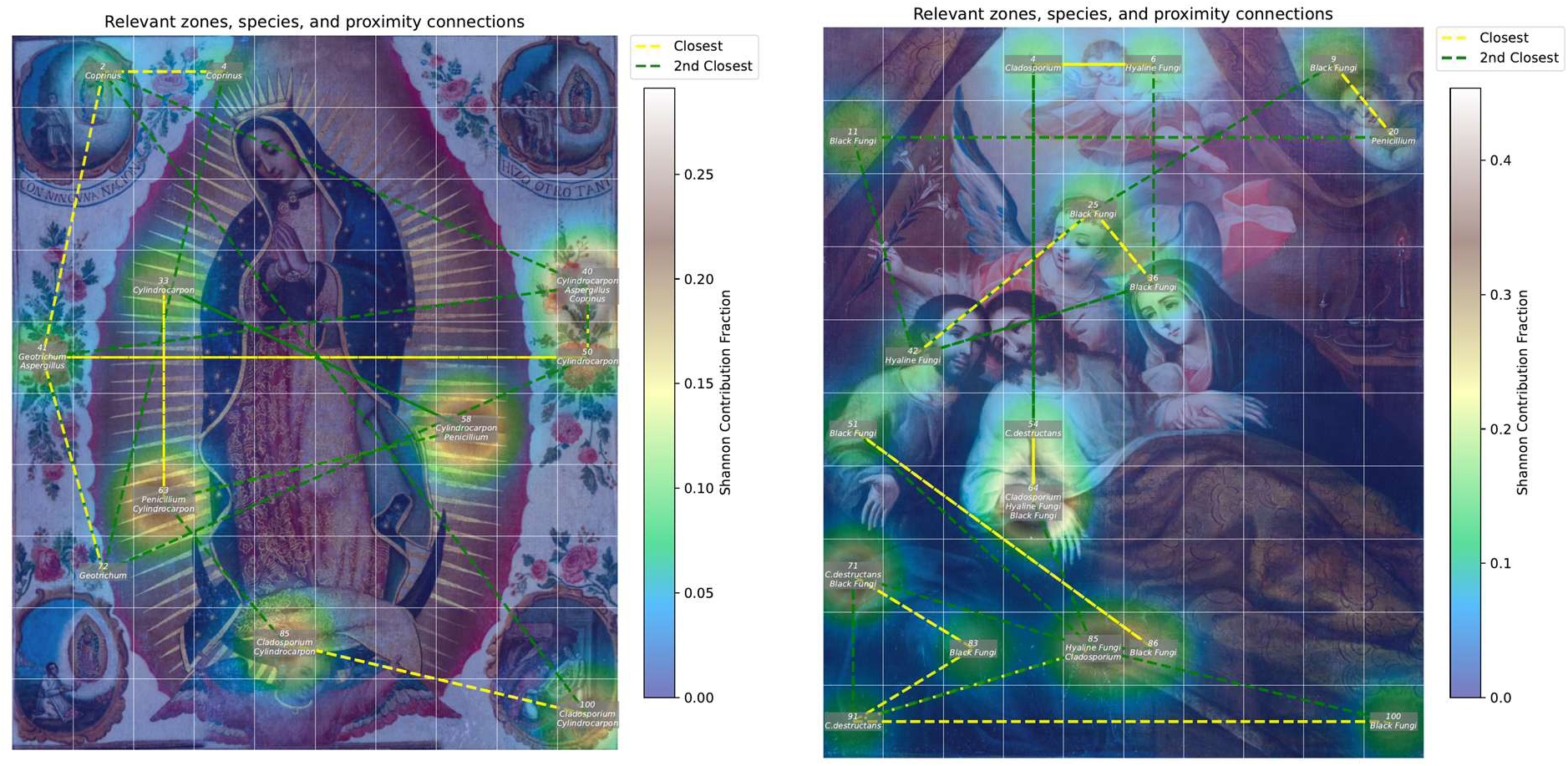
Chromatic proximity, microbial taxa, and Shannon diversity contributions across both artworks. Chromatically similar zones were identified using Bhattacharyya distances in the CIELAB (LAB) color space and are connected by dashed lines (yellow: closest neighbor; green: second-closest neighbor). The background heatmap represents the fractional Shannon diversity contribution of each zone. Overlaid labels indicate the microbial taxa isolated from each corresponding position. (a) *Virgen de Guadalupe*: Several clusters of chromatically proximal zones coincide with repeated isolation of *Cylindrocarpon*, *Penicillium*, and other fungi. (b) *La Muerte de San José*: Spatial convergence between chromatic similarity and microbial distributions is also observed, particularly for black fungi, hyaline fungi, *Cladosporium*, and *C. destructans*.

## Discussion

### Bacterial communities and human-associated taxa

Microorganisms can colonize paintings due to the amount of nutrients and substrates they contain (37). The recovery of multiple *Bacillus* spp. (*B. cereus, B. megaterium, B. altitudinis* and *B. subtilis/amyloliquefaciens/vallismortis*) is consistent with previous reports in which *Bacillus* spp. are among the most frequently isolated bacteria from oil paintings (38). Their capacity to form resistant spores and produce extracellular enzymes and metabolites (3, 39) supports their role in pigment alteration and structural weakening. *B. megaterium* and *B. subtilis* have been previously associated with enzymatic activity and discolored areas in paintings (13), and both were detected in *Virgen de Guadalupe*, suggesting potential involvement in pigment degradation.

The high cellulose content of canvas supports increases their susceptibility to microbial degradation (13). Several genera identified in this study, particularly *Bacillus* sp., are known cellulase producers (40). *B. cereus*, for instance, can synthesize proteases, pectinases, lipases, lectinases and chitinases that may contribute to paint discoloration (23), and has been isolated from cultural heritage documents, where it may cause structural damage (41). Other genera detected, such as *Micrococcus* and *Staphylococcus*, have also been linked to biodeterioration of cultural heritage (42).

Many of the microbial species identified in this study have previously been reported in valuable drawings and manuscripts, including the *Codex Atlanticus* by Leonardo da Vinci, where *M. timonae* and *S. epidermidis* were detected (43). *S. arlettae* has been reported from temples (12), and *Brevibacillus* sp. has been recovered from paint residues and industrial paint waste (44, 45), illustrating the broad ecological range of these taxa.

*M. timonae,* detected in quadrant 98 of *Virgen de Guadalupe*, is noteworthy because species within this genus combine a wide enzymatic repertoire with pigment production (41–44). *M. timonae* has been isolated previously from heritage materials with dark discolorations (41), consistent with its potential role in localized staining of the canvas.

Several isolates corresponded to species associated with human or animal microbiota, including *O. ureolytica, A. lwoffi, S. haemolyticus, S. epidermidis* and *C. firmus* (50–54). In the case of *B. melitensis,* it has been identified as a human pathogen (55). Their presence likely reflects direct human contact and visitor activity, as the paintings are displayed in freely accessible spaces without physical barriers. Although several of these taxa are commonly reported in clinical settings, their detection in this context most likely reflects environmental or human-associated contamination rather than active pathogenic processes.

Species such as *K. rizhophila*, *S. paucimobilis*, and *B. cereus* have previously been reported from canvas paintings in the NTCR (23). *S. paucimobilis* has been associated with museums environments containing paintings and wood (56) and *K. rhizophila* has been documented on historical objects (42). Additionally, bacteria isolated from the wooden frames were dominated by *Bacillus* sp., consistent with reports that these taxa are among the most common wood colonizers, with cellulose-degrading activity that can compromise structural integrity (5, 57, 58).

### Fungal communities and biodeterioration processes

Filamentous fungi are key agents of biodeterioration because their mycelia can penetrate substrates, causing cracking, detachment, color loss, and staining (11). Substrate composition, humidity, water availability, and temperature strongly influence fungal growth; and bacterial biofilms and accumulated dust provide additional nutrients (59). Tropical climates, such as that of Orosi, further enhance these effects (11).

The genera *Aspergillus*, *Fusarium*, *Penicillium* and *Cladosporium,* which were among the most prevalent in both paintings, are ubiquitous and produce abundant airborne conidia. They have frequently been reported in canvas paintings, murals and historic monuments worldwide (11, 37, 59–62). Several species within these genera produce extracellular casein-degrading enzymes and other proteases, enabling degradation of protein-rich substrates such as casein-based binders (63, 64). Canvas supports commonly contain casein, cellulose, proteins, polysaccharides, and oils (3), making them suitable substrates for fungal growth.

*Penicillium* spp., detected in multiple deteriorated and cracked areas (Fig. 2c and Fig. 3c), are well known for causing biodeterioration and inducing chemical changes through enzymatic activity (38). Their high spore production facilitates persistence and recolonization after cleaning (38). Fungi such as *Aspergillus*, *Cladosporium*, *Fusarium* and *Penicillium* also produce pigments (e.g., melanin, quinones, mycosporines, hydroxyanthraquinones, carotenoids) that can generate visible spots and color alterations (62, 63). Some of these pigments have proteolytic and cellulolytic activity (37).

Furthermore, many species of *Aspergillus*, *Cladosporium*, and *Penicillium* produce organic acids capable of dissolving cations and chelating metal ions, leading to the formation of salt crystals that disrupt paint layers and promote cracking and flaking (63). Additionally, the detection of *Pestalotiopsis,* which has shown cellulolytic activity and biodeterioration potential in canvases from the NTCR (65), further underscores the vulnerability of cellulose-based supports.

Additional genera detected in this study, such as *Ulocladium*, *Aspergillus* (various sections), and *Cladosporium*, are characterized by cellulolytic activity (61) and, specifically in the case of *Aspergillus* spp., have been previously associated with deteriorated mural paintings (66). Their presence suggests ongoing degradation of both canvases and wooden elements.

The yeast *T. asahii*, detected in *La Muerte de San José*, is commonly found in tropical environments on decaying wood, soil, air, plants and water, and can also be part of human and animal microbiota (67, 68). Its detection likely reflects a combination of environmental exposure and human presence.

*J. aquatica*, isolated from the surface of *Virgen de Guadalupe*, is typically associated with submerged wood in freshwater ecosystems (63–65) and has been reported from a limited number of regions (72). To our knowledge, this is the first report of *J. aquatica* from painted canvases, highlighting the capacity of heritage surfaces to host unexpected freshwater-associated taxa, possibly introduced via aerosols or splashes. Its specific impact on canvas and paint layers remains to be investigated.

Although *Curvularia* and *Ulocladium* were less frequent, both genera have been associated with biodeterioration in artworks and are favored by high moisture (11, 38). This is particularly relevant in Orosi, which belongs to a rain-rich climatic zone characterized by persistent humidity and high wet-season precipitation (73, 74). Such conditions create favorable microhabitats for fungal growth on cellulose-based materials. *Ulocladium* spp. also possess cellulolytic activity and can degrade cellulose in paper and related substrates (75).

Fungi obtained from the wooden frames, including *Penicillium*, *Aspergillus*, *Fusarium* and Basidiomycota, are typical xylophagous taxa. Basidiomycetes, especially Agaricomycetes, are among the most important wood-decay fungi, capable of degrading cellulose and hemicellulose and modifying lignin (76–78). Their presence indicates active biodeterioration of the frames.

Both paintings were previously restored due to severe deterioration (79). The persistence of bacteria and fungi despite restoration suggests that new materials or retouching layers may provide additional nutrients or microenvironments that support recolonization (80), emphasizing the need for long-term microbiological monitoring.

### Biofilm formation and implications for heritage materials

Biofilms constitute a key survival strategy for bacteria, providing protection against variations in humidity, temperature and pH, while ensuring nutrient distribution and hydration; biofilms typically contain up to 97% water (81). Biofilm formation is a common response to environmental stress and confers increased resilience, metabolic cooperation and environmental stability (82).

On heritage surfaces, biofilms drive biodeterioration through physical and biochemical mechanisms. Physical adhesion and growth can generate mechanical stress, whereas biochemical activity involves secretion of extracellular enzymes and acids that degrade organic and inorganic components, alter pH and ion concentrations, accumulate water and produce pigments (1, 7, 83, 84). These processes weaken materials, accelerate decomposition and increase fragility.

In this study, *S. saprophyticus* and *B. subtilis/amyloliquefaciens/vallismortis* were identified as strong biofilm producers, whereas *K. rhizophila* exhibited moderate biofilm formation (Table 4, Fig. 4). For *S. saprophyticus*, this is consistent with previous reports in which most strains displayed strong biofilm-formation capacity (85). Biofilm formation in staphylococci is frequently mediated by an intercellular polysaccharide adhesin encoded by the *ica* operon (86, 87). Likewise, *B. subtilis* is a widely used model organism for biofilm studies (88) and its biofilms are rich in exopolysaccharides and proteins (89). *K. rhizophila* has also been reported to form highly complex biofilms, particularly in association with other microorganisms (90). The presence of these biofilm-forming taxa on the paintings and frames suggests that microbial communities may be organized in structured consortia capable of sustained colonization and material degradation.

### Essential oil volatiles as candidate biocides

Essential oils have been proposed as alternatives to conventional biocides because they are generally recognized as safe, environmentally friendly and rich in antimicrobial components (18). Their application to cultural heritage has been explored mainly on stone and mural surfaces; their use on canvas paintings is less common due to concerns about possible interactions with organic binding media and pigments (15).

In this study, linalool, α-terpineol, terpineol, eucalyptol, limonene and α-pinene were evaluated against three representative bacterial strains. Linalool was the most effective compound, completely inhibiting growth of all three strains in non-contact assays (Table 5, Fig. 5). Eucalyptol and α-terpineol also showed substantial activity, especially against *B. subtilis/amyloliquefaciens/vallismortis*. These results align with previous work showing that linalool can be more potent than eucalyptol at lower concentrations (16).

Oxygenated terpenoids, such as eucalyptol, linalool, and α-terpineol, generally exhibit greater antimicrobial activity than hydrocarbon terpenes like limonene and α-pinene (16). This pattern was evident here, as limonene and particularly α-pinene were the least active compounds. Among oxygenated terpenoids, the strongest effects were observed for linalool and terpineol. Alcohol-containing terpenoids are lipophilic and can integrate into cell membranes, causing membrane expansion, pore formation, increased fluidity and permeability, disruption of membrane proteins, inhibition of respiration and electron transport, and interference with macromolecule synthesis, ultimately leading to cell death (16).

Similar results were expected for α-terpineol and terpineol (Table 5), given that the terpineol used in this study consists of a mixture of terpineol isomers, among which α-terpineol is typically the predominant component and a major constituent of many essential oils from aromatic plants (91). The observed differences between treatments may therefore reflect experimental variability, as indicated by the standard deviation, or result from antagonistic or synergistic interactions among the various aromatic compounds that comprise essential oils, which can reduce or enhance their antimicrobial activity (92–94).

Differences in susceptibility among bacterial species likely reflect variations in membrane lipid composition, permeability and surface charge (16). In these assays, *B. subtilis/amyloliquefaciens/vallismortis* was the most susceptible, inhibited by all six compounds in most conditions, whereas *S. saprophyticus* was the most resistant.

A key observation was that non-contact assays consistently produced larger inhibition halos than direct-contact assays (average 3.74 cm vs 1.02 cm). This likely reflects the more homogeneous dispersion of volatile compounds in the headspace and the higher dose applied to the lid (100 µL) compared with the filter disk (approximately 10 µL). Because these compounds are relatively hydrophobic, diffusion through the agar matrix in direct contact may be limited, whereas vapor-phase diffusion is enhanced.

Isopropanol, used as a positive control, exhibited lower biocidal activity than most essential oil components and was ineffective in non-contact conditions. In contrast, several components—especially linalool—were active exclusively through their volatile fraction. This observation is significant for conservation practice because the ability to inhibit microbial growth without direct contact with the painted surface reduces the risk of altering pigments or binders and aligns with the principle of minimal intervention (14).

Overall, our results support the potential of selected essential oil volatiles, particularly linalool and α-terpineol, as candidate agents for preventive conservation. However, further work is needed to assess their long-term stability, compatibility with paint layers and varnishes, and efficacy in situ under real museum or church conditions.

### Environmental conditions

A fundamental factor to be considered is the long-term exposure of these artworks to tropical climatic conditions. For example, the interesting thing about the data shown in Fig. 7 is that they can be compared with the values reported for two climatically contrasting sites: the Louvre Museum (approximately temperature of 21 °C and humidity of 40%) (95) and Thailand (roughly temperature of 30 °C, humidity of 60%) (12). Thus, the high relative humidity (12), likely contributed to the elevated microbial diversity observed in these artworks.

### Diversity and Statistical Analysis: comparison with previous tropical artwork studies

The diversity indices obtained for the Orosi paintings showed patterns consistent with those reported for Lanna mural paintings in Northern Thailand (12) (Fig. S5 for detailed indices per artwork). In our study, *La Virgen de Guadalupe* exhibited the highest fungal biodiversity (Chao1 = 24.0; H’ = 2.35; 1–D = 0.87), followed by *La Muerte de San José* (Chao1 = 14.5; H’ = 2.04; 1–D = 0.85).

Suphaphimol *et al.* (12) obtained similar trends between the two Lanna murals with different locations and visitor frequencies. The murals at Buak Krok Luang, characterized by high visitation, exhibited Chao1 ≈ 31.5, H’ ≈ 2.35 and 1-D ≈ 0.85. On the other hand, the murals at Tha Kham, characterized by low visitation, showed Chao1 ≈ 22.5, H’ ≈ 1.75, 1-D ≈ 0.70. Their study identified fungi as a key deteriorative agent on the murals, mainly through production of organic acid compounds and calcium mineral precipitation. These processes cause pigment discoloration and mechanical damage. Further details of microbial richness, diversity, and community dissimilarities across all artworks are presented in the Supplementary Information (Figs. S5–S6). This research, conducted under extreme climatological conditions, contributes to the current understanding of microbiological diversity in two-dimensional artworks. Studies on the Shannon index in three-dimensional and older objects show high values (96, 97); the eighteenth-century paintings examined here show similarly high diversity and are comparable in community complexity.

The patterns observed in Orosi were further contrasted with those reported for indoor-controlled environments at the NTCR (23, 98, 99). As shown in Fig. S5, the *Virgen de Guadalupe* painting exhibited the highest microbial richness and diversity, reflected by elevated Shannon–Wiener and Fisher’s α values, whereas *La Muerte de San José* showed intermediate diversity dominated by a smaller number of taxa. In contrast, samples from the NTCR, including the Management’s Office (99) and *Musas I* (23, 98), displayed lower richness and higher dominance, as indicated by reduced Shannon and increased Simpson indices. These differences suggest that the controlled indoor conditions at the NTCR support simpler and less diverse microbial communities than those developing under the tropical microclimatic conditions of Orosi.

Consistently, Bray–Curtis dissimilarity analysis revealed marked compositional differences between the Orosi and NTCR microbial communities (Fig. S6), highlighting the strong influence of environmental conditions on microbial community structure in cultural heritage settings.

## Concluding remarks

This study revealed a rich microbial community associated with eighteenth-century paintings in Orosi, dominated by genera known to contribute to biodeterioration. Both bacterial and fungal isolates showed enzymatic or biofilm-forming capabilities that can accelerate pigment degradation and canvas deterioration. Environmental data confirmed that the tropical microclimate, characterized by high humidity and temperature, favors microbial colonization, emphasizing the need for region-specific conservation strategies.

Volatile components from essential oils, particularly linalool and α-terpineol, showed strong antibacterial effects without requiring direct contact, suggesting their potential use in preventive conservation. Their volatility allows antimicrobial activity without physical interaction with painted surfaces, aligning with minimal-intervention conservation principles.

Future research should integrate molecular tools for more precise fungal identification and metagenomic approaches to better understand microbial biofilm dynamics on heritage surfaces. Additionally, in situ validation of essential oil volatiles under museum or church conditions is recommended to assess long-term efficacy and material compatibility. Sustainable preservation of tropical heritage artworks will depend on the development of integrated conservation protocols that account for microbial ecology, environmental management, and safe biocide application.

## Acknowledgements

The authors are grateful to Keylin Ureña-Alvarado for preparing the graphical abstract and to Sharon Sandí-Delgado for photographing the paintings. The authors also acknowledge the assistance of Alejandra Gómez-Arrieta in processing the mycological samples, and Laura Villalobos-Soto and Yeimmy Ramírez-Zúñiga for their assistance in the isolation and identification of bacterial samples. The authors thank the Iglesia de Orosi for facilitating access to the artworks.

